# Signalling pathways drive heterogeneity of ground state pluripotency

**DOI:** 10.1101/373373

**Authors:** Kirsten R. McEwen, Sarah Linnett, Harry G. Leitch, Prashant Srivastava, Lara Al-Zouabi, Tien-Chi Huang, Maxime Rotival, Alessandro Sardini, Thalia E. Chan, Sarah Filippi, Michael P.H. Stumpf, Enrico Petretto, Petra Hajkova

## Abstract

Pluripotent stem cells (PSCs) can self-renew indefinitely while maintaining the ability to generate all cell types of the body. This plasticity is proposed to require heterogeneity in gene expression, driving a metastable state which may allow flexible cell fate choices. Contrary to this, naive PSC grown in fully defined ‘2i’ environmental conditions, containing small molecule inhibitors of MEK and GSK3 kinases, show homogenous pluripotency and lineage marker expression. However, here we show that 2i induces greater genome-wide heterogeneity than traditional serum-containing growth environments at the population level across both male and female PSCs. This heterogeneity is dynamic and reversible over time, consistent with a dynamic metastable equilibrium of the pluripotent state. We further show that the 2i environment causes increased heterogeneity in the calcium signalling pathway at both the population and single-cell level. Mechanistically, we identify loss of robustness regulators in the form of negative feedback to the upstream EGF receptor. Our findings advance the current understanding of the plastic nature of the pluripotent state and highlight the role of signalling pathways in the control of transcriptional heterogeneity. Furthermore, our results have critical implications for the current use of kinase inhibitors in the clinic, where inducing heterogeneity may increase the risk of cancer metastasis and drug resistance.

## Introduction

Developmental plasticity is a defining feature of pluripotency and heterogeneity has been proposed as a key regulatory mechanism^1–6^. Pluripotent stem cells (PSCs) can be derived from the pre- and post-implantation embryo, primordial germ cells (the precursors of sperm and oocyte) and via induced reprogramming of somatic cells^1, 7^. These cells can be propagated indefinitely *ex vivo*, while retaining the remarkable capacity to generate all cell lineages of the body either through *ex vivo* differentiation or by re-introducing cells to the *in vivo* environment for embryonic development.

In the mouse pre-implantation embryo, FGF signalling generates a salt-and-pepper pattern of pluripotency and lineage markers such as *Nanog* and *Gata6*^8^. PSCs recapitulate this heterogeneity in the presence of FGF signalling^1, 8–10^. This heterogeneity is dynamic and reversible and such dynamic equilibrium of gene expression in PSCs is defined as metastability^5, 11–14^. Metastability has been proposed as a hallmark of the pluripotent state, safeguarding the downstream capacity for flexible cell fate decisions^1–6^.

Heterogeneity at the single-cell level has been clearly demonstrated in traditional PSC growth environments containing serum^11–14^. The 2i environment is a more recently developed, fully defined medium containing small molecule inhibitors of MEK and GSK3 kinases^15^. The choice of environmental growth condition has a substantial impact on the molecular properties of PSCs, yet cells grown in both conditions remain functionally pluripotent with full developmental potential, demonstrated by their ability to differentiate to all lineages and contribute efficiently to chimeras including the germ line^15–18^. Global gene expression profiles and epigenetic states show a striking difference between serum and 2i environments; DNA methylation is substantially reduced in 2i^18–20^ and histone modifications have altered global levels and genomic distributions^16, 18^. The pluripotent state is thus intrinsically capable of tolerating a flexible molecular profile. Upon switching environmental conditions, molecular profiles are reversed, further emphasising the flexible and dynamic nature of both the transcriptome and epigenome in PSC^16, 18–20^.

Contrary to the PSC heterogeneity detected in serum-containing environments, the 2i condition captures cells in a relatively homogeneous state with respect to pluripotency and lineage markers^10, 16, 21–24^. Inhibition of MEK in 2i prevents the transduction of FGF signals, and thus heterogeneity driven by FGF signalling is lost. This absence of heterogeneity in 2i argues against metastability as a hallmark of pluripotency. In contrast, heterogeneity of the extraembryonic marker *Hex* has been reported in 2i, suggesting the potential for expanded cell lineage potency^25^ and a lengthy unsynchronised cell cycle has been proposed to drive heterogeneity in cell cycle genes in 2i^26^. In view of these findings, it is currently debated whether 2i represents a homogeneous ground state and it remains controversial whether heterogeneity and thus metastability is an inherent feature of pluripotency^1–5, 9, 16, 27–29^. In response to the homogeneity of 2i with respect to pluripotency and differentiation markers, a formative phase linking pluripotency and multilineage progression has been proposed^29^. This intermediate state is proposed to exhibit increased gene expression noise upon release from 2i conditions, allowing differentiation to proceed.

Heterogeneity of PSCs has predominantly been considered in the context of single-cell heterogeneity. However, variation across populations has been well-described in induced PSCs (iPSCs) which show differences across cell lines. This has been attributed to various sources, including genetic variation, epigenetic memory linked to incomplete erasure of the somatic programme, and differences in laboratory protocols^27^. An alternative hypothesis is that heterogeneity is a feature of the pluripotent state, in which case genetically identical PSCs grown under controlled environmental conditions would also show population heterogeneity. Little is known about population heterogeneity across isogenic embryonic stem cell (ESC) lines or across other PSC sources including embryonic germ cells (EGCs) and this may thus represent a currently unknown component of pluripotent metastability.

Here we show by genome-wide transcriptome profiling that 2i conditions cause increased transcriptional heterogeneity at the population level. 2i population heterogeneity is dynamic, supporting the notion of a metastable state of pluripotency. Using this population-based approach we discover that calcium signalling is heterogeneous in 2i and by targeted analysis we demonstrate that this is recapitulated at the single-cell level. Furthermore, we show that 2i kinase inhibition causes loss of signalling robustness regulatory mechanisms via reduced negative feedback to the EGF receptor and we propose this drives the downstream calcium heterogeneity. Overall, we demonstrate that molecular analysis at the population level presents a powerful approach to identify drivers of heterogeneity compared to single-cell studies which, given the inherent technical noise, can accurately quantify only the mostly highly expressed genes^30^.

## Results

### 2i induces population heterogeneity

To assess population heterogeneity in PSCs, we compared 22 isogenic PSC lines grown in traditional serum-containing conditions or in 2i, both containing LIF^18^. We tested two pluripotent cell subtypes, ESCs derived from pre-implantation embryos and EGCs derived from primordial germ cells (see Supplementary Table 1 for details)^7^. Bulk global transcriptomes of all mouse PSC lines were measured and unsupervised dimensionality reduction by multidimensional scaling (MDS) shows that the environmental condition causes a major change to the transcriptome (Fig. 1a, first MDS component), confirming earlier reports^16, 18^. Surprisingly, we also detect increased heterogeneity in 2i relative to serum at the population level for both ESCs and EGCs, measured by sample distance across the second MDS component (Fig. 1a).

**Figure 1.**
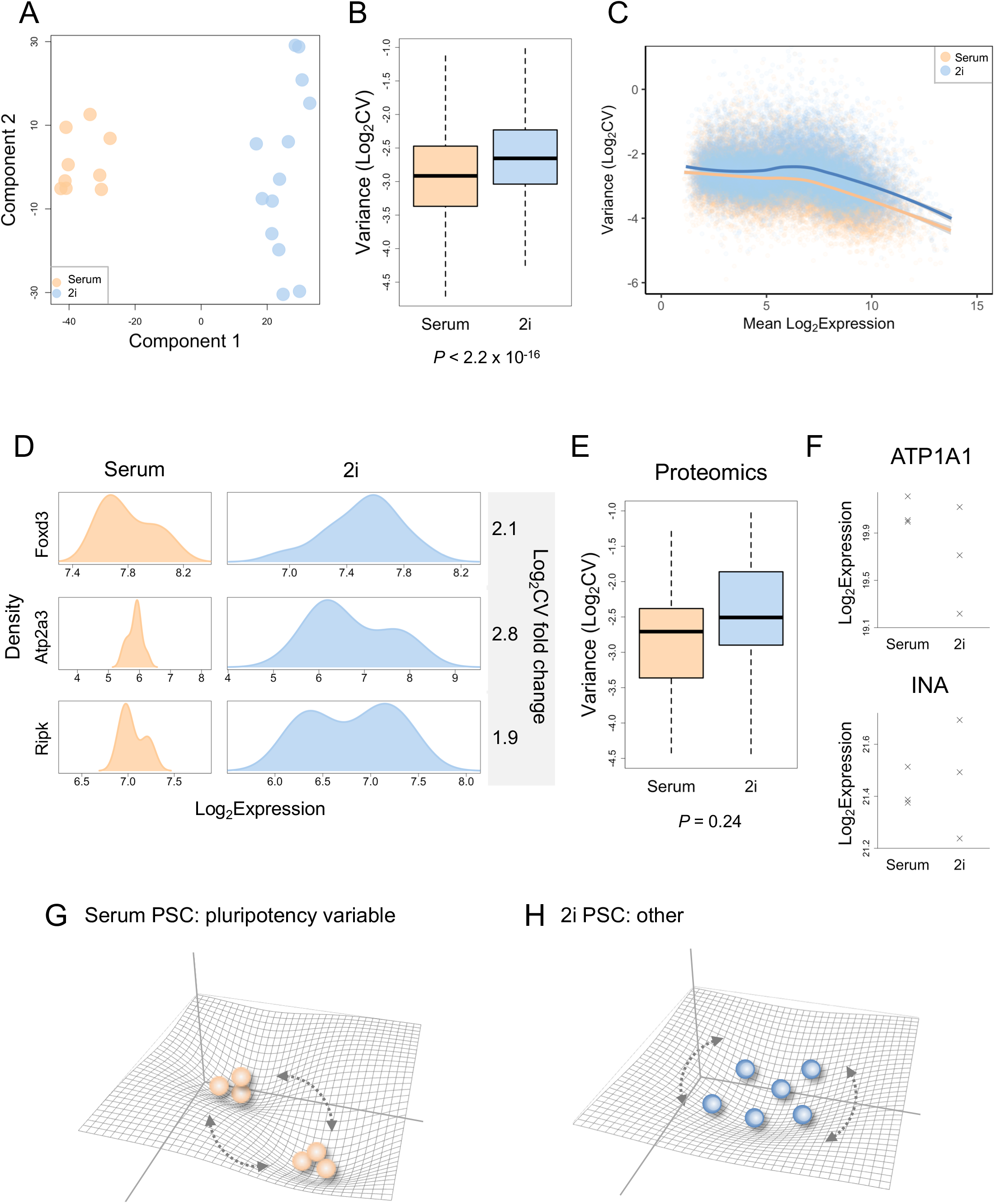
Transcriptional heterogeneity in 2i. **a**, Global transcriptomes were profiled for 22 PSC cell lines in serum and 2i conditions, with each sample representing a population of cells; MDS shows greater heterogeneity in 2i across component 2. **b**, Variance across populations (Log_2_CV) for all genes, showing 2i is significantly more variable than serum; two-sided Wilcoxon signed rank test. **c**, Mean versus variance (Log_2_CV) for all genes depicting greater 2i variance across expression levels. **d**, Density distributions of example genes across populations, with Log_2_CV fold change (2i/Serum). **e**, Variance (Log_2_CV) for proteins of genes with significantly greater 2i heterogeneity relative to serum at the transcriptional level, *n* = 3 biological samples in each condition, two-sided Wilcoxon signed rank test. **f**, Log_2_Expression levels showing greater 2i heterogeneity for example proteins. **g**, Pluripotency genes are heterogeneous in serum at the population level, schematically depicted as balls on a landscape with two attractor states. **h**, Heterogeneous genes in 2i represent a broad attractor state, compared to pluripotency markers in serum which cluster into two states.

To directly quantify population heterogeneity within each environmental condition, we calculated the variance (Log_2_CV) for each gene in 2i and in serum environmental conditions. We identify significantly greater variance in 2i than serum genome-wide (*P* < 2.2×10^−16^, Fig. 1b, Supplementary Fig. 1a,b). The relationship between mean expression levels and variance is the same in serum and 2i, with a trend towards reduced variance for high expressed genes compared to low expressed genes (Fig. 1c), consistent with previous reports^31^. Variance is increased in 2i across the range of mean expression values, with the exception of non-expressed genes (Fig. 1c). This highlights the power of population analyses to detect heterogeneity beyond the most abundant genes. Notably, individual genes show stark differences in variance with a range of distribution profiles (Fig. 1d), in contrast to the small difference observed at the whole-genome level.

To verify that the difference in population heterogeneity between conditions is not due to methodological differences, we performed analysis by qPCR. We tested candidate genes with significantly higher population heterogeneity in 2i (*FDR* < 0.2, Supplementary Table 2). Our results confirm increased heterogeneity in 2i relative to serum at the population level (*P* = 0.016), independently validating our initial findings (Supplementary Fig. 1d).

We next performed proteomics on three serum and three 2i PSC lines. Label-free proteomics quantified the most abundant 1,185 proteins. When sampling the proteins of genes with significantly greater 2i heterogeneity relative to serum at the transcriptional level, there is a trend towards increased heterogeneity in 2i (Fig. 1e). Although we recognise the reduced power to detect any global difference in heterogeneity with small sample sizes (*n* = 3), we nevertheless observe increased heterogeneity in 2i for individual proteins (Fig. 1f), indicating heterogeneity exists at the protein level in 2i.

The preexisting view of pluripotency considers the serum PSC state as heterogeneous with 2i representing a homogeneous ground state. Our observation of greater heterogeneity in 2i suggests otherwise, at least at the population level. Previous studies have shown that serum-cultured PSCs are heterogeneous specifically for pluripotency and lineage markers and we thus sought to determine if pluripotency genes show a different pattern in our data. We find that pluripotency network components maintain reduced heterogeneity in 2i at the population level and are instead more heterogeneous in serum (Supplementary Fig. 1e-g). We note the presence of two pluripotent states in serum as previously reported^26, 32^ and find this is driven by the expression of *Tbx3*, *Nanog* and *Esrrb3* (Supplementary Fig. 1e). We conclude that 2i is homogeneous for pluripotency factors, as reported by others at the single-cell level^10, 16, 21–24^ and shown here at the population level, while other genes show variability in expression in 2i conditions. This argues against 2i as an overtly homogeneous ground state and we schematically depict this heterogeneity as landscape plots in Fig. 1g,h.

### Cell sex contributes to PSC heterogeneity

Cell subtypes frequently underlie heterogeneity and we next sought to test if these cause the difference between serum and 2i population heterogeneity. Such subpopulations may cause differences in mean expression levels between samples, manifesting as population heterogeneity. Regarding ESC and EGC cell subtypes (see Supplementary Table 1), ourselves and others have previously established that only minor differences exist between these in serum and 2i^18, 33^. Nevertheless, to explicitly test if these differences cause greater heterogeneity in 2i, we regressed out the effect of cell subtypes to equilibrate expression between ESCs (*n* = 10) and EGCs (*n* = 12). The significant difference in heterogeneity between 2i and serum remains, indicating these sub-populations are not causing the detected differential heterogeneity (Fig. 2c).

**Figure 2.**
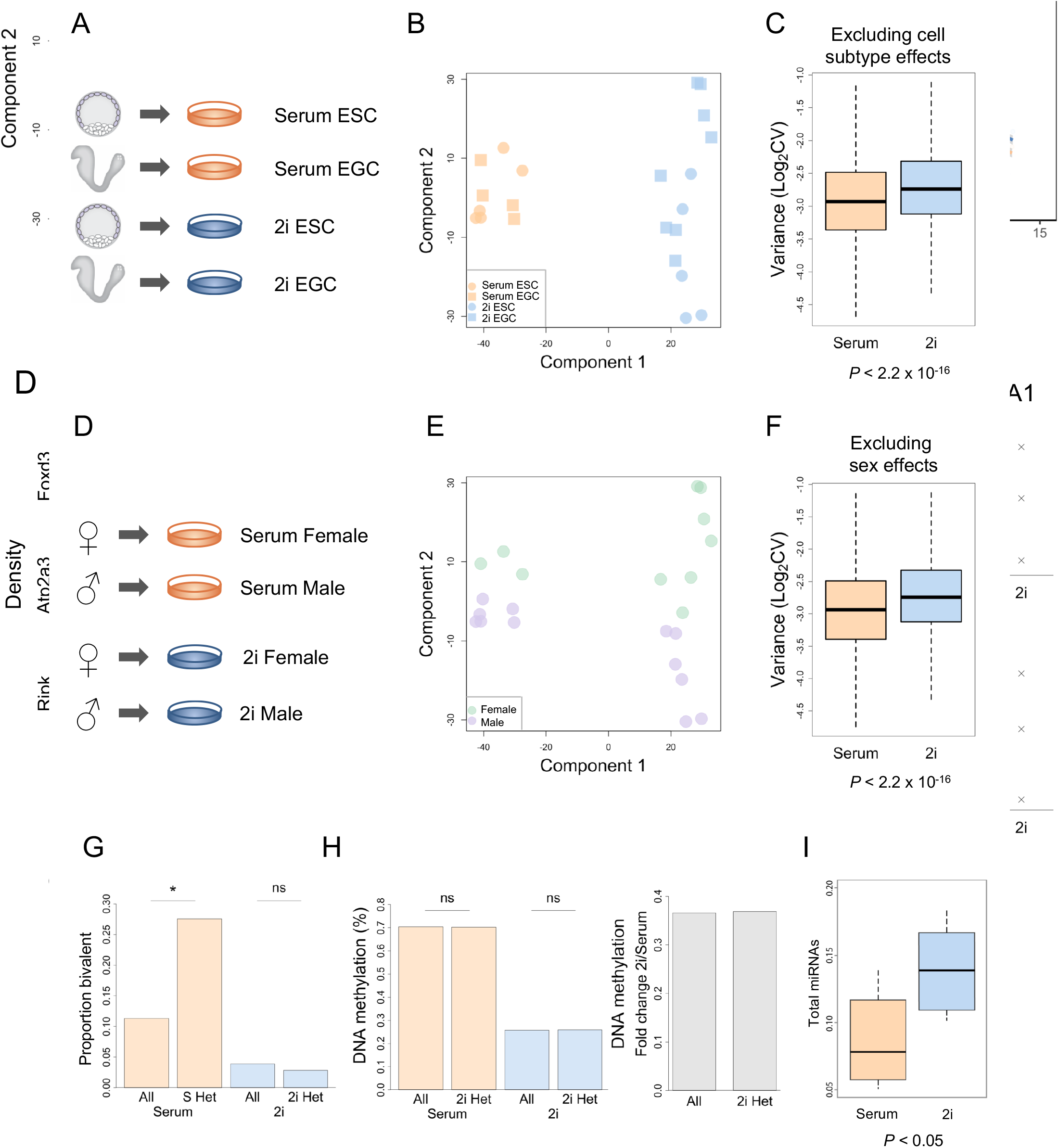
2i heterogeneity is not driven by cell subtypes or epigenetics. **a,** Schematic of ESC (*n* = 10) and EGC (*n* = 12) samples. **b**, MDS of all samples depicting ESC and EGC cell subtypes. **c**, Variance (Log_2_CV) for all genes after regressing out cell subtype showing 2i heterogeneity remains; two-sided Wilcoxon signed rank test. **d**, Schematic of male and female samples, *n* = 10 females, *n* = 12 males. **e**, MDS of all samples showing sex separation across component 2 in serum and 2i. **f**, Variance (Log_2_CV) for all genes after regressing out sex showing 2i-specific heterogeneity is not due to sex differences; two-sided Wilcoxon signed rank test. **g**, Serum heterogeneous genes are enriched for bivalent H3K4me3/H3K27me3 chromatin, 2i heterogeneous genes show no difference to genome average, hypergeometric test. **h,** Gene body DNA methylation is not different for 2i heterogeneous genes compared to the genome average for all genes, including fold change between serum and 2i. **i**, Total miRNA read count, as a proportion of all small RNAs, are not reduced in 2i and are instead increased relative to serum; two-sided Wilcoxon test.

Genetic sex can drive cellular differences^34^ and to test this we examined the transcriptomes of 10 female and 12 male cell lines (Fig. 2d). We identify a clear distinction between female and male samples across the second MDS component; sex therefore underlies the second major transcriptional difference between PSCs after environmental conditions (Fig. 2e). Changes occur to genes involved in signaling (*FDR* = 0.00015, UniProt KW-0597) and metabolism (*FDR* = 0.0073, GO:0006006), consistent with previous reports^33, 35^. Both serum and 2i conditions show differences in gene expression between female and male cells (Fig. 2e), indicating sex subpopulations contribute to part of the observed pluripotent heterogeneity in both environments.

Previous studies show an interaction between sex and environmental conditions in PSCs^35^. To determine if sex differences cause differential transcription heterogeneity between culture conditions, we regressed out the effect of sex and find that the significant difference in culture-heterogeneity remains (Fig. 2f). Thus, sex subpopulations are not driving 2i-specific heterogeneity and instead drive heterogeneity in PSCs regardless of environmental conditions.

### Chromatin bivalency is associated with serum heterogeneity

Bivalent chromatin, consisting of the histone modifications H3K4me3 and H3K27me3, has recently been associated with transcriptional heterogeneity in PSCs^22, 36^. These studies tested serum PSCs, for which bivalency is more common across the genome than in 2i (Fig. 2g, ‘All’)^16^. We show that genes with greater 2i heterogeneity are not enriched for bivalent chromatin compared to the genome average (*P* > 0.05, Fig. 2g, ‘2i Het’). This is in contrast to the few genes we detect more heterogenous in serum at the population level (*n* = 127), for which bivalency is significantly enriched (*P* < 0.05, Fig. 2g, ‘S Het’). This lack of association between chromatin bivalency and 2i-specific heterogeneity suggests distinct regulatory mechanisms for these genes compared to serum-specific heterogeneous genes.

Heterogeneity has also been proposed to be regulated by gene body DNA methylation^4, 24, 37–39^. DNA methylation is substantially reduced under 2i conditions^18–20^ and we hypothesised that this may drive the difference in transcriptional heterogeneity between environmental conditions. We find no difference in gene body DNA methylation for 2i heterogeneous genes in comparison to all genes and show the fold change between serum and 2i is unaffected (Fig. 2h). Thus, the hypomethylated state of 2i is not associated with increased transcriptional heterogeneity for these genes.

We tested additional regulatory features that differ between serum and 2i, including miRNA biogenesis and RNA polymerase II travelling ratio^16^. *Dicer* and *Drosha* components of the miRNA biogenesis machinery are downregulated at the transcriptional level in 2i (Supplementary Fig. 1h) and a reduction in total miRNA abundance may cause decreased transcriptional robustness. However, we show that total miRNAs, as a proportion of all small RNAs, are not reduced and are instead increased in 2i (Fig. 2i) arguing against this. This may be a consequence of increased expression of *Argonaute* biogenesis machinery components (Supplementary Fig. 1h), or due to post-transcriptional regulation of miRNA synthesis. We also find that the RNA polymerase II travelling ratio is not significantly altered between serum and 2i for genes with greater 2i heterogeneity (gene set enrichment analysis (GSEA), *FDR* > 0.05, Supplementary Fig. 1i).

### Heterogeneity reflects dynamic gene expression

The 2i environment enables derivation of PSCs from strains and species previously resistant to *ex vivo* culture^7^. 2i may thus possess the ability to capture a broader range of stable pluripotent states than serum growth conditions. In such a case, expression differences between cell lines, representing stable pluripotent subtypes established during the derivation procedure, would cause population heterogeneity. We therefore profiled three different cell lines derived in 2i and tracked these over time to see if each cell line has a distinct transcriptome state. The same procedure was undertaken in serum conditions. All six cell lines were grown in parallel, using the same media batch for 2i or serum, and global transcriptomes were profiled at passage 11, 14 and 17 (Fig. 3a). The results of our initial experiment are reproduced, with greater sample distance in 2i across the second MDS component (Fig. 3b). Variance is significantly increased in 2i (*P* < 2.2×10^−16^, Fig. 3c), across the range of expressed genes (Supplementary Fig. 2a).

**Figure 3.**
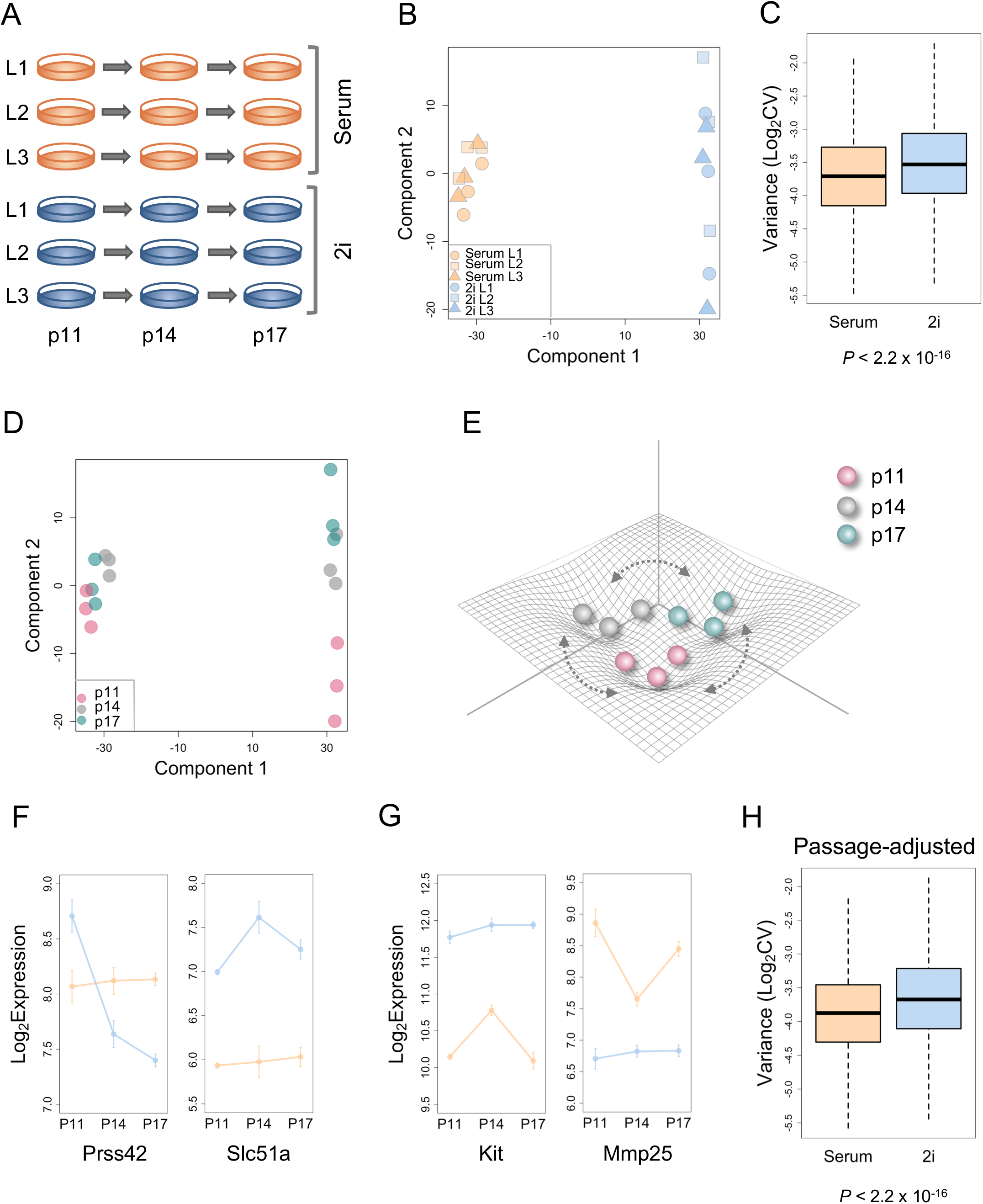
Heterogeneity reflects dynamic gene expression. **a**, Experimental design for transcriptome-wide profiling over time. EGC lines: L1, L2, L3; Passage: p11, p14, p17. **b**, MDS of all samples, reproducing greater heterogeneity in 2i across component 2. Cell lines (L1, line 1; L2, line 2; L3, line3) do not cluster together indicating heterogeneity is not due to intrinsic, stable differences between cell lines. **c**, Variance (Log_2_CV) across all genes is significantly higher in 2i; two-sided Wilcoxon signed rank test. **d**, MDS showing global gene expression changes over time in serum and 2i. **e**, Schematic landscape of metastability over time in both serum and 2i. **f**, Dynamic and reversible gene expression across passages in 2i and **g**, in serum. **h**, Variance (Log_2_CV) for all genes after regressing out passage indicating dynamic gene expression is not driving heterogeneity specifically in 2i; two-sided Wilcoxon signed rank test.

Heterogeneity that arises via the capture of distinct stable states during each derivation process in 2i is expected to cause the clustering of samples by cell line, with the transcriptome of each cell line remaining constant over time (Supplementary Fig. 2b). Instead, we show that each cell line changes its global expression profile over time, with samples at a given time point more closely related to each other than each cell line (compare Fig. 3b with Fig. 3d,e). This occurs in both serum and 2i environments.

Heterogeneity is thus not due to intrinsic differences between cell lines but is malleable and dynamic in both serum and 2i.

A range of dynamic expression profiles is detected, including genes stable in serum and dynamic in 2i (e.g. *Prss42* and *Slc51a*, Fig. 3f) and vice versa (e.g. *Kit* and *Mmp25*, Fig. 3g). These expression fluctuations are reversible and exemplify dynamic, metastable gene expression in PSCs under both environments. Furthermore, heterogeneity is not amplified over time, but fluctuates with passage (Supplementary Fig. 2c,d). Of note, dynamic gene expression over time is a greater contributor to the transcriptome than sex, as indicated by the loss of clustering of female and male samples across MDS component 2 which is replaced by passage clustering (Supplementary Fig. 2e). We confirm 2i heterogeneity is not driven by fluctuations to MEK and GSK3 inhibitor levels induced by experimental variation during culture, as target genes of these pathways do not have increased heterogeneity in 2i (Supplementary Fig. 3a). Overall, our results show PSCs are metastable at the population level regardless of growth environment, genetic variation and laboratory conditions. This is also independent of stable intrinsic differences between cell lines and we further show metastability does not bias individual cell lines towards a particular developmental lineage (Supplementary Fig. 3b). Expression of developmental regulators are not upregulated over time and pluripotency factor gene expression remains high (Supplementary Fig. 1g, Supplementary Fig. 3c,d), showing maintenance of the pluripotent state in both conditions and the absence of lineage bias with passage. Overall, the detection of dynamic heterogeneity in both serum and 2i PSCs suggests that this metastability over time may be a feature of pluripotency.

To test if dynamic gene expression causes the difference in heterogeneity between serum and 2i, we regressed out passage. 2i remains more heterogeneous than serum after removing the passage effect, indicating that while dynamic expression is a source of PSC heterogeneity, it does not drive the serum-2i difference (Fig. 3h). We further show that 2i heterogeneity remains after regressing out cell subtype, sex and passage effects (Supplementary Fig. 3e).

### 2i causes calcium signalling pathway heterogeneity

To identify the underlying drivers of 2i-specific population heterogeneity, we explored the expression and functions of genes with significant differential variance between environmental conditions (*FDR* < 0.2, *n* = 280, 2i > serum, Supplementary Fig. 4a). Fold change ranges from 1.7 to 5.9 (Fig. 4a) and expression levels range from low to high (Supplementary Fig. 4b). Functional ontology analysis^40^ shows that these genes are enriched for phosphoprotein and endoplasmic reticulum genes, pointing toward heterogeneity in signalling pathways (*FDR* < 0.05, Fig. 4b). 2i inhibits MEK and GSK3 signalling molecules and we confirm that the downstream target genes of these pathways^35^ are significantly repressed as expected (Supplementary Fig. 4c), while heterogeneity is not increased in 2i (Supplementary Fig. 3a). To test other downstream signal pathway target genes in an unbiased manner, we searched for transcription factor motifs^41^ that are enriched at the promoters of 2i heterogeneous genes (+/− 200 bp from the transcription start site). This revealed a striking enrichment of CREB (calcium/cAMP response element binding protein) and ATF (activating transcription factor) motifs (*P* < 5 × 10^−4^, Fig. 4b), implying that 2i heterogeneous genes are regulated by these transcription factors.

**Figure 4.**
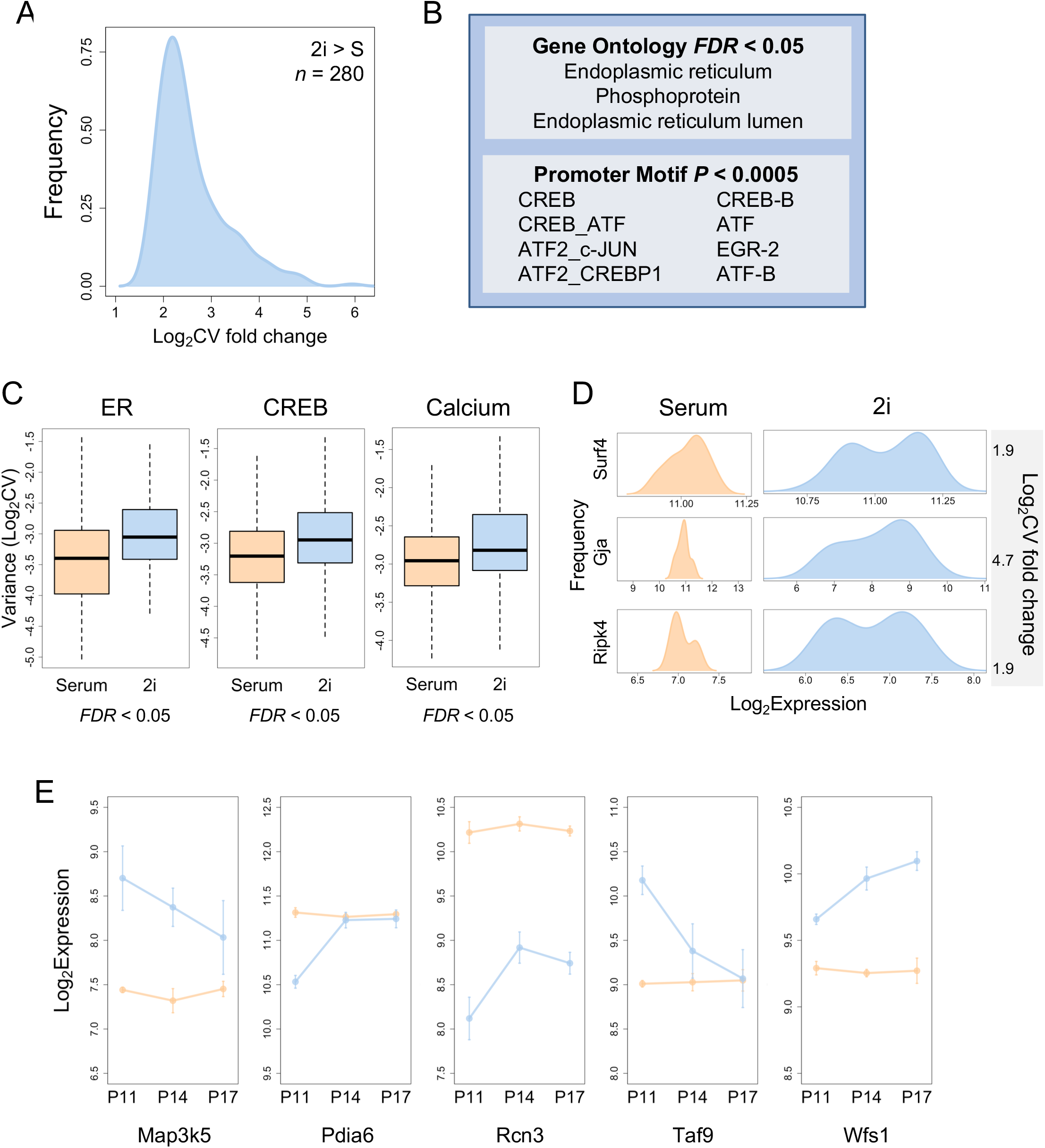
Signalling pathways are heterogeneous in 2i. **a**, Fold change in variance (Log_2_CV) density distribution for genes with significantly greater heterogeneity in 2i than serum (*FDR* < 0.2, *n* = 280). **b**, Gene ontology and promoter motif enrichment for genes with significantly greater heterogeneity in 2i. **c**, Variance (Log_2_CV) is increased in 2i beyond the global level for endoplasmic reticulum (ER, *n* = 166), CREB target genes (*n* = 257) and calcium signalling (*n* = 181); statistics performed against genome average using GSEA (*FDR* < 0.05, NES > 1.3). **d**, Density distributions of gene expression within enriched gene ontology groups across population samples. **e**, Dynamic expression of calcium, endoplasmic reticulum and CREB target genes in 2i compared to serum.

The CREB/ATF family of transcription factors show concerted regulation by many pathways, with canonical regulation via calcium signalling. The endoplasmic reticulum, identified by gene ontology analysis, is a key regulator of calcium signalling and acts as an intracellular calcium ion store. We thus explicitly tested the variance of the entire calcium signalling pathway (Kyoto Encyclopedia of Genes and Genomes (KEGG), *n* = 181)^42^. We find that calcium signalling heterogeneity is significantly higher in 2i than serum, beyond the global level (GSEA *FDR* < 0.05, normalised enrichment score (NES) > 1.3, Fig. 4c). Affected genes include calcium binding proteins, membrane bound proteins and transcription factors with a range of distribution profiles (Fig. 4d). We also confirm the entire set of endoplasmic reticulum (KEGG, *n* = 166)^42^ and CREB target genes (Salk Institute, *n* = 257)^43^ show the same response (GSEA *FDR* < 0.05, NES > 1.3, Fig. 4c). Furthermore, calcium, endoplasmic reticulum and CREB target genes are dynamic, with gene expression fluctuating over time (Fig. 4e).

Thus, alteration of signalling pathway states through constitutive inhibition of MEK and GSK3 converges on increased calcium signalling heterogeneity.

### Calcium is heterogeneous at the single-cell level in 2i

Population data reflects the conglomerate gene expression of single cells and previous reports have suggested population heterogeneity can be used as a proxy for single-cell heterogeneity^44–46^. The pluripotent transcription factor *Nanog* exhibits greater heterogeneity in serum than 2i at the single-cell level^11, 47, 48^ and we show this also occurs at the population level (Supplementary Fig. 1g), indicating this gene shows the same pattern in both single-cell and population data.

To experimentally test if population heterogeneity can be directly used to predict single-cell heterogeneity in 2i, we next quantified transcription at the single-cell level (Fig. 5a). Candidate genes with significantly higher population heterogeneity in 2i were tested by qPCR across a range of functional pathways, including calcium, endoplasmic reticulum and CREB target genes, general transcription factors and WNT and AKT pluripotent signalling pathway genes (Supplementary Table 2). Mean expression levels correlate well between single-cell and population data (*Pearson cor* > 0.91, Supplementary Fig. 5a,b).

**Figure 5.**
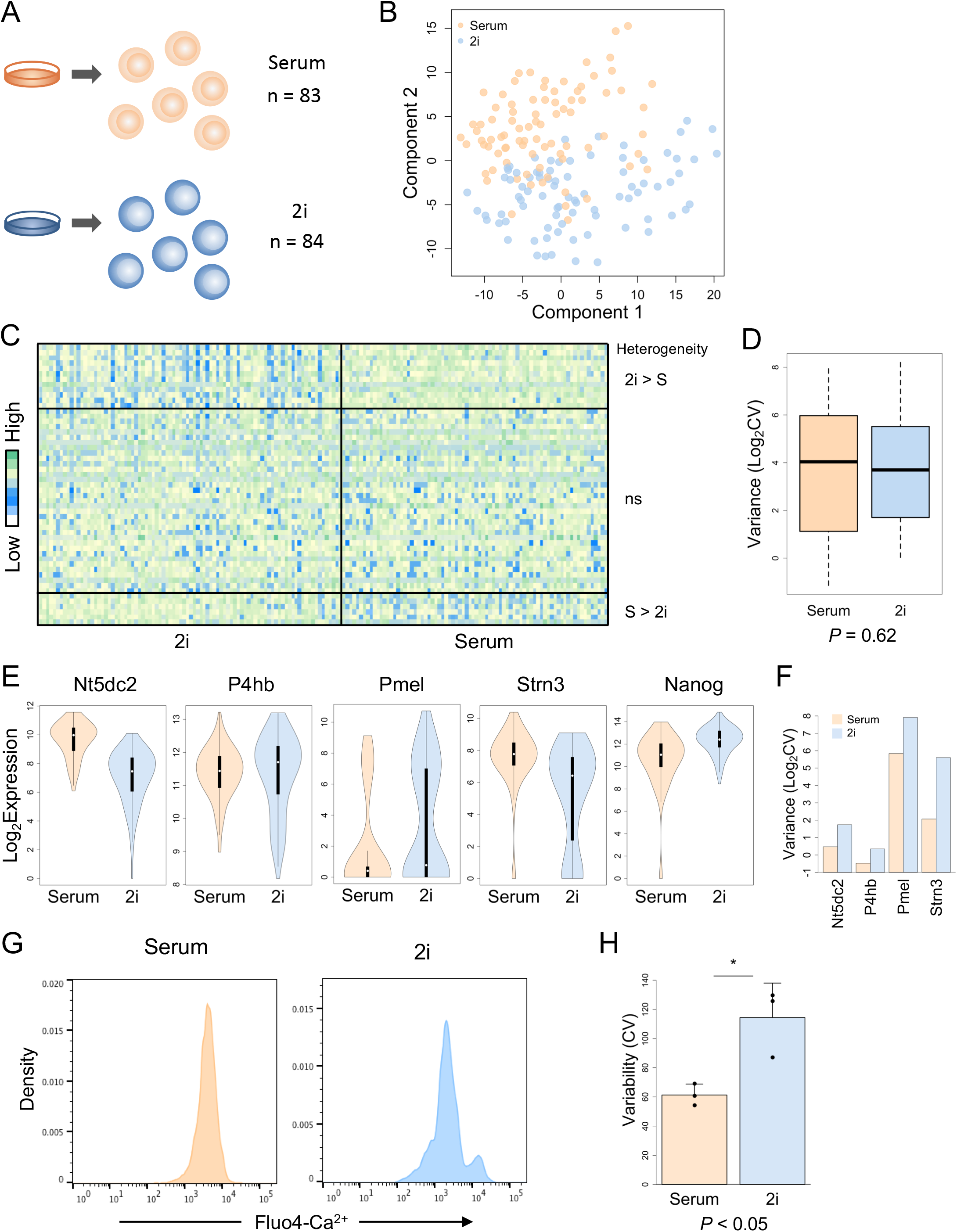
Single-cell calcium signalling heterogeneity in 2i. **a,** Schematic of single-cell transcription experimental design. **b,** MDS of all samples in serum and 2i (*n* = 47 genes). **c**, Heatmap of single-cell gene expression showing increased variability in 2i compared to serum for a subset of genes (upper panel, *FDR* < 0.05) and a large proportion of genes with a non-significant difference between 2i and serum (ns, *FDR* > 0.05, middle panel). Genes with increased serum heterogeneity relative to 2i from previously published reports were also included (lower panel, *Esrrb*, *Hes1*, *Klf4*, *Nanog*, *Nr0b1*, *Tbx3*). **d**, Variance (Log_2_CV) does not differ across all genes; two-sided Wilcoxon signed rank test. **e**, Single-cell qPCR violin plots for genes with significantly greater heterogeneity at the single-cell level in 2i (*FDR* < 0.05). *Nanog* is heterogeneous in serum as expected. **f**, Variance (Log_2_CV) for example genes. **g**, Heterogeneity of intracellular calcium messenger levels is higher in 2i than serum, measured by flow cytometry of Fluo4-Ca^2+^ fluorescent indicator. Representative of 3 biological replicates. **h**, Flow cytometry variance of intracellular calcium levels from 3 biological replicates.

Of particular importance, we identify novel genes with greater single-cell transcriptional heterogeneity in 2i compared to serum (*FDR* < 0.05, Fig. 5c upper panel, Fig. 5e,f, Supplementary Fig. 5c), proving single-cell heterogeneity indeed exists in 2i conditions. As demonstrated by heatmap, a subset of genes show increased variability in gene expression in 2i, with more homogeneous expression in serum (Fig. 5c upper panel). Distributions of gene expression and quantification of the variance (Log_2_CV) are depicted for example genes in Fig. 5e,f. However, for the majority of genes tested, no significant difference in variance was detected between environmental conditions (Fig. 5b-d, Supplementary Fig. 5d). These results indicate that not all heterogeneity observed at the population level is directly replicated at the single-cell level. Overall, our results show that population heterogeneity cannot be used as a general proxy for single-cell heterogeneity. Genes that do show consistent heterogeneity between single-cell and population levels are specific to calcium, endoplasmic reticulum and CREB targets, suggesting that functional pathways are a more accurate predictor of single-cell heterogeneity than the individual gene level. In further support of our data, analysis of independent single-cell datasets from Teichmann and colleagues confirm the 2i heterogeneous calcium signalling genes identified by population analysis have increased transcriptional heterogeneity in 2i relative to serum PSC conditions at the single-cell level (*P* < 0.05, one-sided Wilcoxon signed rank test)^26^.

To test if calcium messenger signalling is affected at the single-cell level beyond transcription, we quantified intracellular calcium levels by flow cytometry to measure single-cell variance. We show heterogeneity of calcium signalling is clearly impacted by 2i (Fig. 5g). A significant increase in heterogeneity is detected relative to serum, reproduced over three biological experiments (*P* < 0.05, Fig. 5h). Overall, our findings show the heterogeneity initially detected by global transcriptome profiling and confirmed at the single-cell level impacts messenger signalling in 2i PSCs and thus that heterogeneity exists in PSCs independent of either growth environment, albeit in different pathways.

### 2i disrupts regulators of signalling robustness

Cellular signalling is robustly regulated to maintain homeostasis and feedback loops are a well-established mechanism reducing heterogeneity^49^. MEK signalling leads to the activation of downstream negative feedback regulators, including dual-specificity phosphatase, protein phosphatase and sprouty families^50^. This MEK-induced negative feedback has been shown to confer robustness^50–54^ and we thus hypothesised that MEK inhibition in 2i leads to a loss of negative feedback that drives heterogeneity.

Our results show that negative feedback regulators downstream of MEK are significantly reduced upon 2i signal inhibition at both the population level (GSEA *FDR* < 0.05, NES > 1.3, Fig. 6a, Supplementary Fig. 6a,b) and single-cell level (Supplementary Fig. 6b). These genes are among the most strongly downregulated genes in 2i, with over 10-fold reduction in expression relative to serum. The EGF receptor is a known target of MEK-induced negative feedback^55^ and has a dual role in activating both MEK signalling and the release of intracellular calcium stores from the endoplasmic reticulum. In 2i conditions, we show negative feedback to the EGF receptor is significantly reduced. Phosphorylation is lost at the Thr693 target site (Fig. 6b,c, Supplementary Fig. 6c), which normally inactivates the receptor in response to MEK signalling.

**Figure 6.**
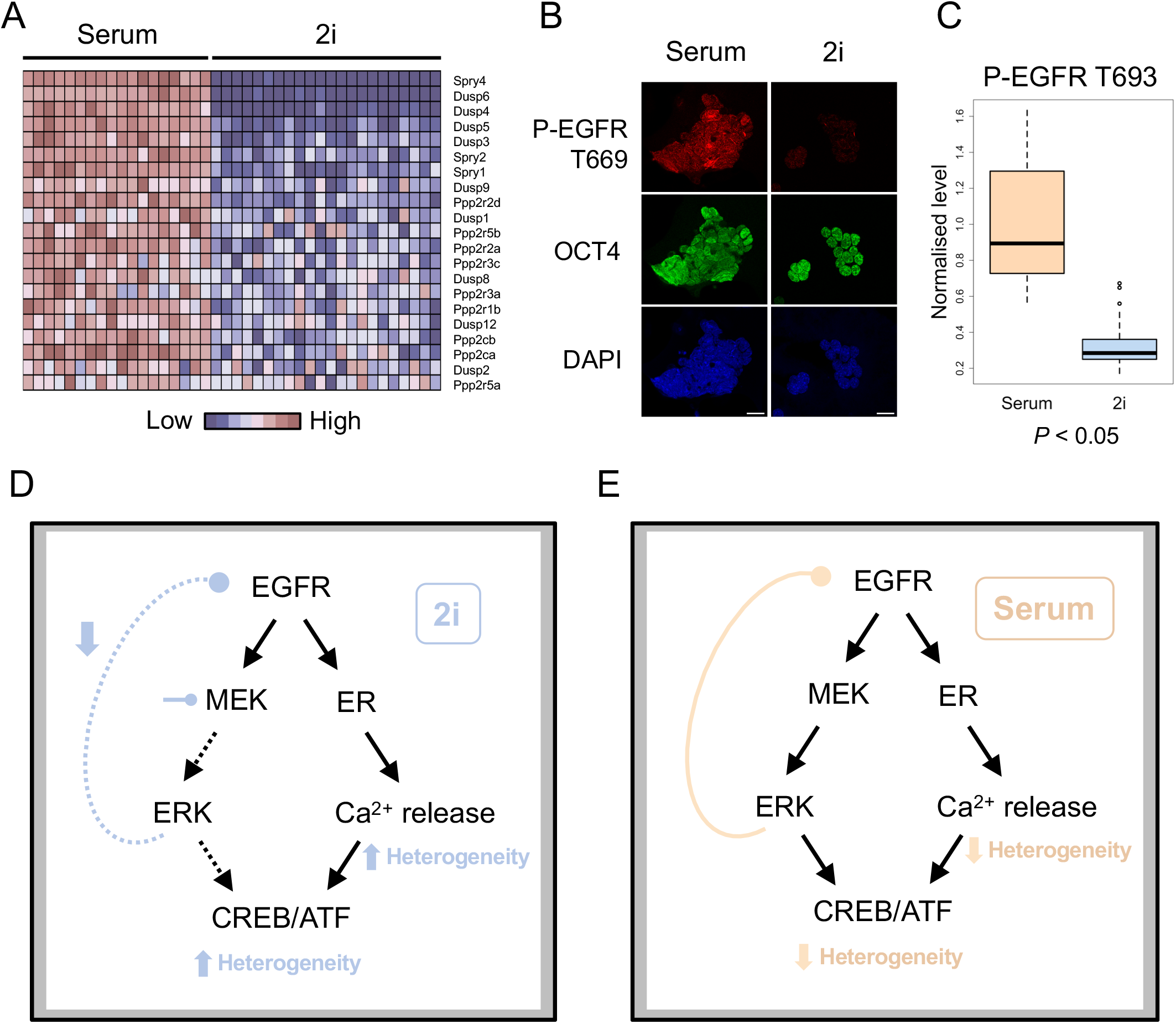
Negative feedback is reduced in 2i. **a**, Heatmap of population level gene expression for negative feedback regulators reduced in 2i. **b**, Immunofluorescence showing the negative feedback site P-EGFR T693 is reduced in 2i. P-EGFR T693, red; OCT4, green; DAPI, blue pseudocolours. Representative of three technical replicates and two biological replicates. **c**, Quantification of P-EGFR T669 immunofluorescence negative feedback. Boxplot outliers defined as points; *n* = 30 cells. **d**, Mechanistic model for increased heterogeneity driven by MEK inhibition in the 2i environment compared to **e**, serum conditions. In 2i conditions, MEK inhibition reduces ERK signalling, causing loss of negative feedback to the upstream EGF receptor. Relief of this negative feedback loop, which is an established regulator of robustness, consequently drives heterogeneity in the parallel calcium signalling pathway.

These results suggest MEK inhibition reduces negative feedback to the upstream EGF receptor in 2i conditions and in line with the well-established role for negative feedback in regulating robustness^49–54^, we propose this causes downstream heterogeneity in the parallel calcium signalling pathway (Fig. 6d,e).

## Discussion

Here we demonstrate the power of using population transcriptomics to identify drivers of heterogeneity in stem cells. Heterogeneity at the population level has previously been observed across distinct iPSC lines, however this has been attributed to diverse genetic backgrounds and incomplete erasure of epigenetic memory^27^. We compare isogenic PSCs grown under controlled laboratory conditions and our results, surprisingly, uncover increased population heterogeneity in 2i compared to serum growth environments. We show that this heterogeneity is not due to stable pluripotent sub-states arising during derivation, but is dynamic and fluctuates over time.

We further show that heterogeneity exists in 2i at both the population and single-cell level in the calcium signalling pathway, caused by constitutive kinase inhibition. MEK inhibition reduces FGF signalling in PSCs, which normally drives heterogeneous expression of pluripotency and lineage markers^1, 8–10^. In the absence of FGF signalling, we confirm these genes are homogenously expressed in 2i, in agreement with previous reports^10, 16, 21–24^. We show that 2i signal inhibition concurrently reduces known regulators of signalling robustness that are downstream of the MEK signalling pathway. Negative feedback regulators induced by MEK normally constrain signalling heterogeneity^50–54^ and their reduction in 2i culminates in reduced negative feedback to the EGF receptor and consequently leads to increased heterogeneity in the parallel downstream calcium signalling pathway (see model diagram in Fig. 6).

Heterogeneity of pluripotency and lineage markers in serum PSCs has been shown to affect the propensity of cells to differentiate^11, 48, 56^. This heterogeneity is dynamic, for example *Nanog*-low and *Nanog*-high cells maintain the capacity to reverse expression and contribute to all somatic lineages^11^. This metastable state is proposed to allow flexible cell fate decisions, reflecting a hallmark of pluripotency^1–6, 9, 16, 28^. Our results present a paradigm shift of the ground state of 2i PSCs; heterogeneity in signalling pathways proves heterogeneity is not limited to serum conditions and occurs in PSCs regardless of growth environment. The specific components of each environmental condition lead to distinct targets, with serum driving heterogeneity in FGF target genes and the signalling inhibitors in 2i bypassing the FGF signalling pathway and driving calcium heterogeneity.

PSC plasticity is marked by dynamic gene expression, an adaptable transcription factor circuitry and an interchangeable epigenetic state^4, 16, 18–20, 29, 57, 58^. DNA methylation is rapidly reversible in PSCs^18–20^ and core histones are hyperdynamic^59^. This plasticity coincides with the potential to form any cell type of the body. Furthermore, the ability to adapt is inherent to cells of the early embryo, which can reverse cell fate upon heterotopic grafting and can tolerate cell biopsy^60, 61^. The dynamic equilibrium of gene expression observed in PSCs indicates a metastable state, where cells are able to explore the state space without loss of pluripotency^5, 11–14^. Moreover, we find that time-course analysis of the steady state (Fig. 3d) reproduces snapshots of gene expression (Fig. 1a), suggesting PSCs show hallmarks of ergodicity in which the system behaves the same over time as over snapshots of the state space^5^. Such metastability may provide the scope for flexible cell fates and experimental evidence is now emerging suggesting heterogeneity precedes developmental decisions^62–67^. Our findings highlight the critical function of signalling pathways, which are the leading determinants of developmental decisions, in the control of transcriptional heterogeneity.

The single-cell and population heterogeneity that arises through kinase inhibition is especially relevant for cancer. Kinase inhibition is a primary therapeutic strategy and our finding that transcriptional heterogeneity is induced by constitutive kinase inhibition has profound implications for such therapies. Stem cells share parallel features with cancer cells^68^ and current inhibitor therapies may have unintended consequences of driving both inter- and intra-tumour heterogeneity. Heterogeneity can drive tumour development across a broad spectrum of cancers, presenting opportunities for drug resistant subclones and metastatic potential to arise^69^. Urgent attention is required to test if clinically in-use MEK inhibitor therapies in melanoma and non-small cell lung cancer impact heterogeneity and consequently impact the risk of drug resistance and metastasis.

Our results underline the need for a systems level understanding of signalling pathway architecture and regulation in the context of heterogeneity. This will improve the design of targeted therapies for regenerating, directing differentiation or destroying stem and cancer cells to improve patient therapies.

## Methods

### Animal studies

Animal studies were authorized under a UK Home Office Project License and carried out in a Home Office-designated facility.

### Cell culture

ESCs and EGCs (Oct4ΔPE-GFP) were derived and cultured as previously described^18^. Serum culture conditions contain 15% FCS, 0.1 mM MEM nonessential amino acids, 2 mM l-glutamine, 1 mM sodium pyruvate, 0.1 mM 2-mercaptoethanol and 1000U/mL LIF in DMEM-F12 with maintenance on a mouse embryonic fibroblast (MEF) feeder layer. 2i culture conditions contain 1 μM PD0325901, 3 μM CHIR99021 and 1000U/mL LIF in N2B27 medium, maintained on laminin (10 μg/ml). Cells were incubated at 5% CO_2_, 95% relative humidity. All samples were profiled at fewer than 23 passages. For time-course transcriptome profiling EGCs were derived and maintained in either serum or 2i and collected every 3 passages from 11-17.

### Population transcriptomics

Samples were prepared and processed as described previously. Briefly, 100ng RNA was fragmented, labelled and hybridized to Affymetrix GeneChip Mouse Gene 1.0 ST Arrays. Data was preprocessed in R using robust multiarray averaging, variance stabilising transformation and Combat batch correction^70^. Limma was used for differential expression analysis and for linear regression against cell subtypes, sex and passage^71^. Differential variance was tested using the studentised Breusch-Pagan test with culture condition as the independent variable. Testing statistical enrichment of gene sets was performed using functional annotation ontology^40^, gene set enrichment analysis (GSEA)^72^ and the TRanscription factor Affinity Prediction (TRAP) tool for enrichment of promoter transcription factor motifs^41^. Boxplot elements are defined by center line, median; limits, upper and lower quartiles; whiskers, 1.5x interquartile range, outliers excluded.

### Proteomics

Proteomics was performed on 12 samples including 3 biological replicates of each condition and 2 technical replicates. Cells were washed twice with ice-cold PBS before resuspension in 8M urea, 20mM HEBES buffer. Samples were vortexed then sonicated with three 10 sec pulses at 20% amplitude prior to centrifugation at 20,000 g for 10 min. Supernatants were collected and Liquid Chromatography-Mass Spectrometry performed at the MRC London Institute of Medical Sciences Proteomics Facility. Analysis was performed with MaxQuant using *FDR* < 0.01 for protein identification and including only those present in at least 10 samples.

### Real time qPCR

Reverse transcription was performed using random-primed Invitrogen Superscript III on 1 μg RNA from population samples. qPCR cycling (70°C 40min, 60°C 30s, 95°C 60s, 30 cycles of 96°C for 5s, 60°C for 20s, followed by a melting curve from 60-95°C in 1°C increments) was performed on 3 μL of 1:40 diluted cDNA. Genes with significantly greater 2i heterogeneity relative to serum were sampled from microarray data and primers with <80% efficiency were removed (*n* = 38 genes tested). 8 biological replicates (*n* = 4 ESC and *n* = 4 EGC) with 3 technical replicates each were tested for serum and 2i. No template and no reverse transcriptase controls were negative.

### Epigenetics

DNA methylation for genes with greater 2i heterogeneity was assessed across gene bodies in serum (*n* = 3) and 2i (*n* = 3). Genome-wide data was sourced from Ficz et al., 2013^19^. Chromatin-immunoprecipitation for H3K4me3 and H3K27me3 was assessed for promoters (2Kb) of 2i-specific heterogeneous genes and serum-specific heterogeneous genes and compared to the genome-wide average^16^.

### Small RNA sequencing

Small RNA transcript cDNA libraries were prepared using the NEBNext Multiplex Small RNA Library Prep Set for Illumina. Briefly, total RNA was extracted using Trizol and 1 μg RNA was used to prepare the cDNA library, including adaptor ligation and reverse transcription.

For multiplexing, 12 sets of index primers were used for 12 cycles of PCR. Amplified cDNA constructs were subjected to size selection using AMPure XP Beads. cDNA ranging from 140-150bp was selected and sequenced by Hiseq2000 (50bp, single end).

### Single-cell transcription

The Fluidigm C1 system with the ClonTech SMART-Seq v4 protocol was used with medium 10μm-17μm integrated fluidic circuits (IFCs) using the redesigned 2016 IFCs to avoid high rates of stacked doublet cells occurring with earlier models (Fluidigm PN 101-3328 B1 white paper). ERCC RNA spikes were added at 1:10,000 to lysis buffer for 12G4 ESCs and amplification was performed at 20 cycles. Single cell capture was confirmed by microscopy and picogreen dsDNA quantification before qPCR on the Fluidigm Biomark HD system using SsoFast EvaGreen (70°C 40min, 60°C 30s, 95°C 60s, 30 cycles of 96°C for 5s, 60°C for 20s, followed by a melting curve from 60-95°C in 1°C increments). Primers were designed spanning an intron and within 2 Kb of the transcript termination site when possible, with product size limited to 80-120 bp. Technical reproducibility was tested during optimisation, median technical CV was 0.85% and therefore only biological replicates were used for downstream experiments. qPCR efficiency for all primer pairs was >80%. No template and no reverse transcriptase controls were negative in all experiments. Poor quality samples and low expressed genes were removed^73^. 47 genes with significantly greater 2i population heterogeneity were quantified. Log_2_Ex was calculated by subtracting the limit of detection of 25 cycles^73^, batch correction was performed^74^ and differential variance was tested using the Brown-Forsythe test.

### Immunofluorescence

Cells were grown on Lab-Tek slides, washed twice with PBS and fixed in 2% PFA for 20min at room temperature. Three PBS washes were performed before permeabilisation and blocking in staining buffer (10% horse serum, 0.3% Triton X-100 in PBS) for 1hr at room temperature. Primary antibody was incubated in staining buffer at 4°C overnight and washed three times with staining buffer before secondary antibody incubation at room temperature for 1hr. Cells were washed in staining buffer once and in PBS twice, before DAPI staining, rinsing with PBS and slide mounting with Vectashield. Cells were imaged using a Leica SP5 confocal microscope. EGFR-T693 antibody: Bioworld BS4063, 1:50 dilution.

### Flow cytometry

Calcium was measured in live E14 ESCs with Fluo4-AM labelled calcium indicator. ESCs were dissociated and MEFs were removed by serial panning for serum cultured cells. 2 × 10^5^ cells were incubated with 3mM Fluo4 and 1.5mM probenicid for 30min at 37°C, 5% CO_2_. Cells were washed twice in 1.5mM calcium-balanced culture medium containing 10mM HEPES then immediately measured by flow cytometry. Ionomycin (10μM) was added at the end of the experiment to confirm cell dye loading. Cells were gated by forward scatter/side scatter (FSC/SSC) followed by FSC-A/FSC-H gating to exclude cellular aggregates. Negative boundaries were determined using Fluo4-AM unloaded cells.

## Data availability

Data are deposited in the Gene Expression Omnibus (GEO) under accession number GSE116565.

## Acknowledgements

We thank members of the McEwen and Hajkova laboratories for stimulating discussions, M. Charalambous for comments on the manuscript and the MRC London Institute of Medical Sciences Flow Cytometry, Microscopy, Proteomics and Genomics Facilities for support. This work was funded by an Imperial College London Fellowship to K.R.M. and Medical Research Council funding to P.H., who is a recipient of the ERC CoG “dynamicmodifications” and a member of the EMBO Young Investigator Programme.

## Author Contributions

K.R.M. and P.H. conceived the study and K.R.M. led the project. Experiments were performed by K.R.M., S.L., H.G.L, L.A-Z., A.S. and T.C.H. Analysis was performed by K.R.M., P.S. and T.E.C.; M.R. and S.F. provided statistical advice. K.R.M. wrote the manuscript with help from P.H. and all authors reviewed and approved the manuscript. Funding was provided by K.R.M. and P.H. for consumables and by K.R.M., M.P.H.S., E.P. and P.H. to support students and staff, including funding to K.R.M to establish her independent research programme.

